# A New Bioink for Improved 3D Bioprinting of Bone-Like Constructs

**DOI:** 10.1101/2021.11.04.467312

**Authors:** Adam C. Marsh, Ehsanul Hoque Apu, Marcus Bunn, Christopher H. Contag, Nureddin Ashammakhi, Xanthippi Chatzistavrou

## Abstract

Bone tissue loss can occur due to disease, trauma or following surgery, in each case treatment involving the use of bone grafts or biomaterials is usually required. Recent development of three-dimensional (3D) bioprinting (3DBP) has enabled the printing of customized bone substitutes. Bioinks used for bone 3DBP employ various particulate phases such as ceramic and bioactive glass particles embedded in the bioink creating a composite. When composite bioinks are used for 3DBP based on extrusion, particles are heterogeneously distributed causing damage to cells due to stresses created during flow in the matrix of the composite. Therefore, the objective of this study was to develop cell-friendly osteopromotive bioink mitigating the risk of cell damage due to the flow of particles. Towards this end, we have linked organic and inorganic components, gelatin methacryloyl (GelMA) and Ag-doped bioactive glass (Ag-BaG), to produce a hybrid material, GelMA-Ag-BaG (GAB). The distribution of the elements present in the Ag-BaG in the resulting hybrid GAB structure was examined. Rheological properties of the resulting hydrogel and its printability, as well as the degree of swelling and degradation over time, were also evaluated. GAB was compared to GelMA alone and GelMA-Ag-BaG nanocomposites. Results showed the superiority of the hybrid GAB bioink in terms of homogenous distribution of the elements in the structure, rheological properties, printability, and degradation profiles. Accordingly, this new bioink represents a major advance for bone 3DBP.

## INTRODUCTION

A major clinical challenge in orthopedic and craniomaxillofacial (CMF) surgery is the development of critically sized bone defects typically caused by congenital malformations, trauma, infection, cancer, or surgical resection [1, 2]. Multiple treatment approaches have been implemented to address such defects such as the use of allographs [3], however, the gold standard continues to be the use of autografts [4]. Globally, over two million bone graft procedures are performed annually [5, 6] with the commonality of such procedures being second only to blood transfusions [7]. The use of autografts is, however, limited by availability, donor-site morbidity [8], and the challenges of creating the required shape (e.g. CMF applications) [9]. While the use of allografts can address the resource limitations and eliminate concerns regarding donor-site morbidity associated with the use of autografts, the use of allografts presents its unique challenges. For example, the use of devitalized allographs not only employs a high-cost laborious process [7], but also results in limited revascularization [10] significantly increasing the risk of incurring an immunological reaction [11] and infection [12]. Treatment strategies for addressing infected bone often require a second revision surgery that not only prolongs recovery time but also has the potential to lead to more severe ramifications such as permanent loss of function and even amputation [13]. Therefore, there is a critical need to develop effective alternative approaches.

To address this, biomaterials have been used, classically as acellular constructs with a focus on identifying biocompatible and bioinert materials [14]. Success has been demonstrated in addressing the shortcomings of traditional biomaterials-based strategies by tackling said challenges through more biomimetic approaches. The advantages of using cell-seeded polymeric tissue engineering constructs were demonstrated in various studies [15–23]. Unfortunately, many of the available polymers are not cell-friendly due to their synthetic origins. Improvements to the cell-friendly nature of biomaterials have been realized through the use of naturally-derived biomaterials such as decellularized tissue matrices [24] or natural polymers such as collagen [25] and gelatin [26]) either incorporated in conjunction with other biomaterials such as synthetic polymers and growth factors or used alone. The enhanced cell-friendly characteristics were in part due to the presence of the tripeptide arginyl-glycyl-aspartic acid (RGD) peptide sequences [27] in said naturally derived biomaterials known to be an important factor in cell attachment and function. While acellular biomaterial-based approaches have represented important advances in the biomaterials field, the inclusion of cells appears to be critical to further advancements. Furthermore, the use of cell seeding alone has proven to be an inadequate approach given the subsequent premature failure of implants due to inhomogeneous cell distributions.

Three-dimensional printing (3DP) technologies allowed significant advancements to be made in controlling the geometry of tissue engineering. Combined with imaging and design technologies such as computed tomography (CT), magnetic resonance imaging (MRI), and computer-aided design (CAD) [28], 3DP enabled the development of customized bone substitutes. A crucial innovation in the expansion of 3DP technologies for tissue engineering was the advent of 3D bioprinting (3DBP) allowing cell-laden constructs to be 3D printed leading to advances in the engineering of biomimetic living constructs [29–31]. Three-dimensionally bioprinted patient-specific constructs show potential for improving treatment outcomes and accelerating the time of recovery. This technology is expected to revolutionize the biomaterials and biomedical engineering fields with such innovations starting to enter clinical studies [32, 33].

The creation of a printable bioink is achieved through the incorporation of cells into a liquid matrix that undergoes solidification post-print, forming the desired tissue-like constructs [34, 35]. Most commonly, 3DBP relies on extrusion-based methods for the printing process [36] however such methods are known to reduce cell viability due to the introduction of shear forces during extrusion. The incorporation of solid elements into bioinks such as glass or ceramic particles that may support bone growth, osteopromotive, leads to increased shear forces during 3DBP leading to additional decreases in cell viability [37].

Typically, bioinks used for 3DBP of bone-like constructs incorporate osteopromotive elements [38, 39] such as Ca-based bioceramics [40, 41] or silicate-based biomaterials [42, 43] in the form of either micro-sized [43] or nano-sized particles [31, 35, 37, 41, 42]. The particles are introduced into the cell-laden polymer matrix through mixing allowing composites to be 3DBP [44]. For example, nanoparticles (NPs) of silicate glasses have been combined with gelatin methacryloyl (GelMA) to form nanocomposites with the two components held together by ionic interactions [45]. Such nanocomposite bioinks are limited to the degree of homogeneity they can achieve not only by the size of the particles incorporated but also by the degree of agglomeration [37, 46]. This was evidenced by the increase in dead osteoblasts after the extrusion 3DBP of bioactive glass (BaG) containing cell-laden bioinks as a result of the increased shear forces [37] introduced by the BaG particles [37]. Therefore, alternative strategies to incorporating osteopromotive elements need to be investigated.

An innovative approach to overcoming these limitations is to chemically link the osteopromotive component(s) to the polymer matrix. It is expected that the chemical bonding of the osteopromotive component(s) and polymer matrix will result in a stable structure allowing agglomeration of osteopromotive component(s) to be minimized. Furthermore, to achieve the greatest degree of homogenization between the osteopromotive component(s) and the polymer matrix will likely require an *in situ* synthesis method. Combining these factors should deliver 3DBP bone-like constructs, where the body cannot distinguish the individual components used in the bioink. This would allow for advanced material characteristics to be achieved that could not otherwise have been realized using a composites approach.

BaG is an attractive osteopromotive component for incorporation into cell-laden bioinks for 3DBP of bone-like constructs given its well-documented improvements in cell viability, osteogenic differentiation, antibacterial properties, and *in vivo* cell survival, as well as anti-inflammatory, and pro-angiogenic properties [47–53]. The ideal BaG-containing bioink, therefore, is likely to exhibit the following characteristics: (1) supports the optimal osteogenic response for cells to be induced as a result of the ions provided by the BaG [50, 51], (2) increases the stiffness of the 3DBP bone-like constructs [52], possesses the necessary anti-inflammatory characteristics for healing, and (4) allows for the promotion of angiogenesis.

We, therefore, aimed to utilize an innovative approach that combined GelMA with an Ag-doped bioactive glass (GAB) to deliver a novel osteopromotive and antibacterial material. To achieve this, we used a methacryloyl functionalized collagen-derived gelatin (GelMA) as the polymer matrix to both enable crosslinking and enhance cell attachment and function. An Ag-doped bioactive glass (Ag-BaG) was selected for its osteogenic, angiogenic [38, 44, 54], and antibacterial properties [55–57], along with the ability of the Ag-BaG to enhance the strength of the delivered material. The GelMA and Ag-BaG were then chemically linked to produce the hybrid, GAB. Additionally, Ag-BaG NPs were mixed with GelMA to deliver a nanocomposite material, where our developed GAB hybrid material was found to be superior to the synthesized nanocomposite material demonstrating the effectiveness of our innovative approach.

## 1. MATERIALS & METHODS

### 2.1. Material synthesis

To synthesize GelMA, type A gelatin (300 bloom; Millipore Sigma) was dissolved in dimethyl sulfoxide (DMSO; Millipore sigma) at 50°C and stirred at 500 RPM. Methacrylic anhydride (MAA; Millipore Sigma) was added in 1/6^th^ increments every 30 minutes to achieve a final MAA: lysine ratio of 2.2 [58]. The resulting solution was then added to toluene at 3x the reaction volume used to induce precipitation of the GelMA [59]. The solution was then decanted after 24h, washed thrice with distilled water, and then dissolved in distilled water at 50°C. After dissolution, the solution was placed at 37°C for 24h to keep the GelMA dissolved while allowing for any remaining toluene to evaporate. The solution was then frozen and lyophilized before storage.

To synthesize the GAB hybrid material, the lyophilized GelMA was dissolved at 10% (w/v) in DMSO at 50°C and stirred at 500 RPM. A coupling agent, (3-Glycidyloxypropyl)trimethoxysilane (GPTMS; Millipore Sigma), was then added to the solution to achieve a hydroxylysine, lysine, arginine to GPTMS ratio of 2.00 and allowed to react for 24h to ensure sufficient time for the epoxy ring-opening reaction (**Fig. 1**). The sol-gel process was used to synthesize the Ag-BaG following previously described methods [56]. 3% (w/w) of the Ag-BaG sol was added to the GelMA solution in addition to distilled water [56] to ensure sufficient hydrolysis between the GPTMS and Ag-BaG components and allowed to stir for 24h to achieve a homogenous distribution between GelMA and Ag-BaG at the molecular level (**Fig. 1**). This solution was then precipitated, washed, and lyophilized as previously described to prevent the materials characteristics from changing during storage (**Fig. 1**). Prior to further use, lyophilized GAB was dissolved at 10% (w/v) in 1X phosphate buffered saline (PBS) along with the photoinitiator VA-086 at 1.5% (w/v) and photopolymerized at 385 nm for 120 s producing GAB hydrogels (**Fig. 1**).

**Figure 1.**
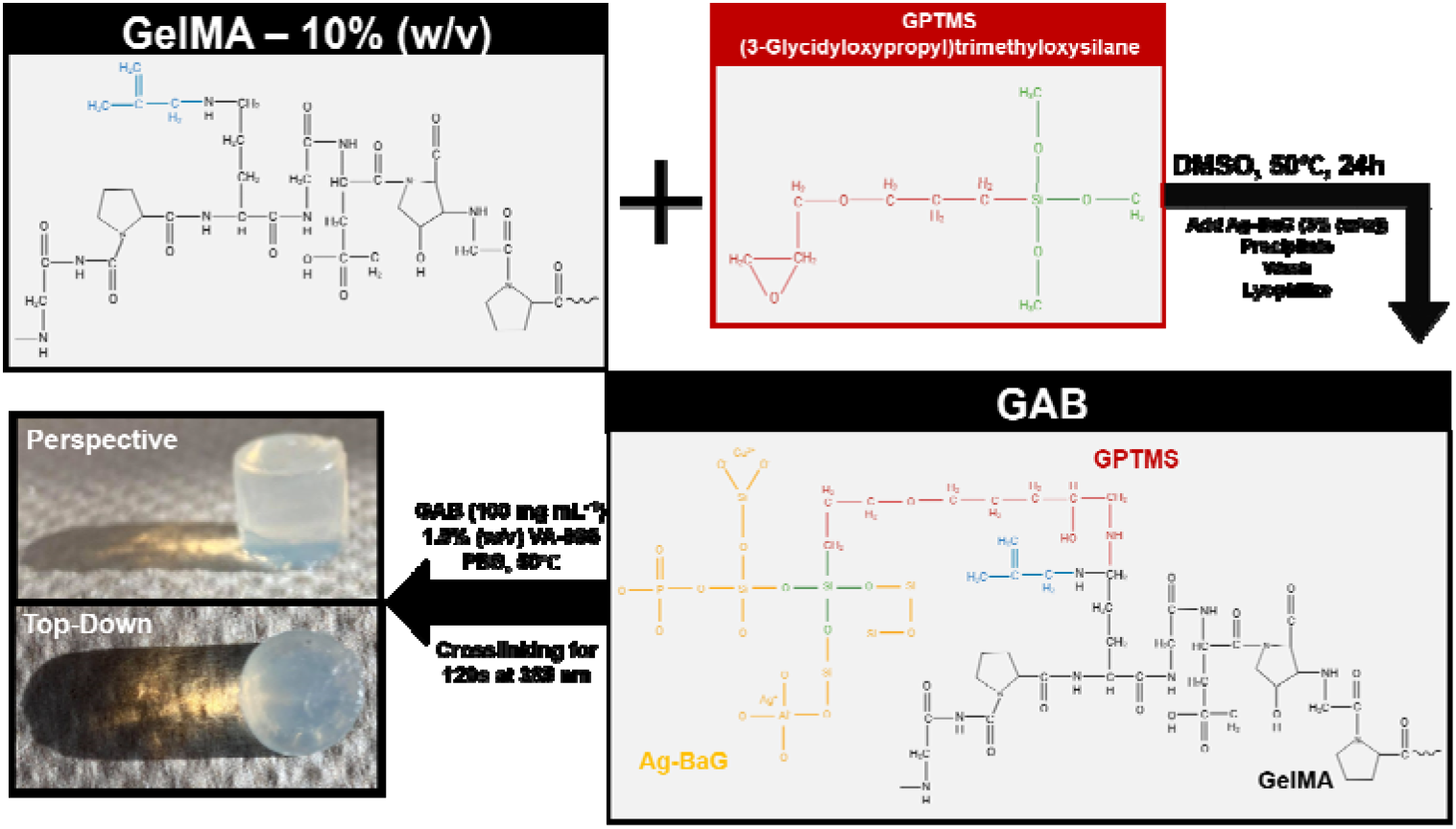
Molecular schematic of the synthesis used to produce GAB along with the process applied to deliver GAB hydrogels.

### 2.2. Structural characterization

The overall morphological characteristics of the synthesized materials were evaluated using optical microscopy (VHX-600E Digital Microscope) and micro-computerized tomography (Micro-CT; Rigaku Quantum GX).

To study the microscopic morphological features of the synthesized materials, scanning electron microscopy (SEM; Tescan MIRA/JEOL 6610LV) was used. Samples were prepared for SEM examination by first undergoing a graded series of ethanol dehydration (i.e. 2x – 25%, 2x – 50%, 2x – 75%, 2x – 90%, and 3x 100% ethanol). The samples were then critically point dried using liquid CO_2_ in order to preserve the native structure of the materials before being metalized with Os_(g)_ for 15s to prevent a buildup of a negative electrical charge. Sample morphologies were captured using a beam voltage of 5 kV. Energy dispersive spectroscopy (EDS; Ametek EDAX Apollo X) was additionally performed to assess elemental homogeneity using a beam voltage of 20 kV.

To identify the molecular bonds present within the synthesized samples, Fourier-transformed infrared spectroscopy – attenuated total reflection (FTIR; Jasco FT/IR-4600) was applied, where spectra were collected from 4000 – 400 cm^−1^. Additionally, spectra relating to the reagents used during the synthesis of GelMA and GAB were collected in order to utilize FTIR for quality control.

Proton nuclear magnetic resonance (^1^H – NMR) was used to determine the methacrylation of gelatin yielded after performing the GelMA synthesis [60]. The 1D spectra were collected using an Agilent DirectDrive2 500 MHz NMR spectrometer, where a sample concentration of 50 mg mL^−1^ in deuterium oxide (D_2_O) was used for all measurements. The degree of substitution (DS) was calculated using the lysine integral method, where the integral of the lysine methylene signals of the gelatin as-received was compared to the integral of the lysine methylene signals in GelMA, as described in equation 1 [61].

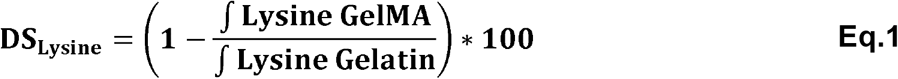

### 2.3. Performance characterization

To assess the swelling behavior of the synthesized samples, lyophilized samples were measured to quantify the mass of the lyophilized samples. The lyophilized samples were hydrated in 1X PBS forming samples 8 mm in diameter having a thickness of 1.5 mm and kept in solution for 24h at 37°C. The hydrated samples were removed from the solution and excess PBS was gently blotted from the surface of the samples. The swelling ratio of the samples was calculated using equation 2.

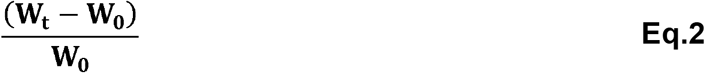

In equation 2, W_t_ represents the weight of the swollen sample after 24h of immersion in PBS, and W_0_ represents the weight of the lyophilized (dry) sample.

Samples were immersed in 1X PBS at 37°C under shaking at 175 RPM to study their degradation behavior. After 1, 3, 5, 7, 14, 21, and 28d of immersion, the pH was measured, and extracts were collected at each time point to elucidate the ion release profile by inductively coupled plasma – optical emission spectrometer (ICP-OES). The samples at each time point were weighted to determine the mass loss profile and determine the time point at which complete degradation is expected.

The viscoelastic behavior of the samples was evaluated using rheological means. To this end, the storage (G’) and loss modulus (G”) of the samples was evaluated as a function of temperature ranging from ambient conditions to 40°C. The viscosity of the samples was evaluated at 37°C using a shear rate from 0.1 to 100 s^−1^.

The compressive behavior of the samples was evaluated using fully cross-linked samples. Samples 8 mm in diameter having a thickness of 1.5 mm were used and the compression testing was performed using a United SFM electromechanical series universal testing machine with a 20 N load cell. A constant cross-head speed of 0.5 mm s^−1^ was used and all samples were compressed up to 60% strain. The elastic modulus was determined by the slope of the linear curve in the elastic region.

### 2.4. Printability

The printability of the samples was investigated by varying the pressure (psi), needle gauge, extruder temperature (°C), speed (mm s^−1^), layer height (mm), cross-linking time (s), and cross-linking intensity (%) during 3D printing (Allevi 3). Semi-quantification of the printability was performed according to the previously reported procedures [62]. In brief, the circularity of the printed enclosed area was calculated using equation 3.

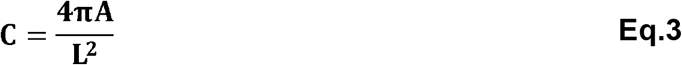

In equation 3, L represents the perimeter, A represents the area, and C represents the circularity, where a circularity of 1 represents a perfectly circular entity. Given the circularity for squared shapes is equal to 1/4A, the printability parameter (P_r_), which is a quantification of similarity of the nature of square-formed prints, was calculated using equation 4.

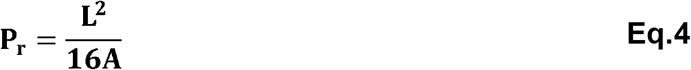

The optimal printability is achieved when P_r_ = 1 and denotes that the ideal viscosity during the print was achieved. It is important to note, as well, that when P_r_ > 1, the viscosity during the print was too high, and P_r_ < 1 denoting the viscosity was too low during the print.

## 2. RESULTS & DISCUSSION

**Figure 2a** showed the ^1^H – NMR spectra of the as-received gelatin, the synthesized GelMA, and the GAB. Using **equation 1**, the degree of substitution for GelMA and GAB using the lysine integral method was found to be ~100% providing supporting evidence that the GAB synthesis did not compromise the degree of substitution.

**Figure 2.**
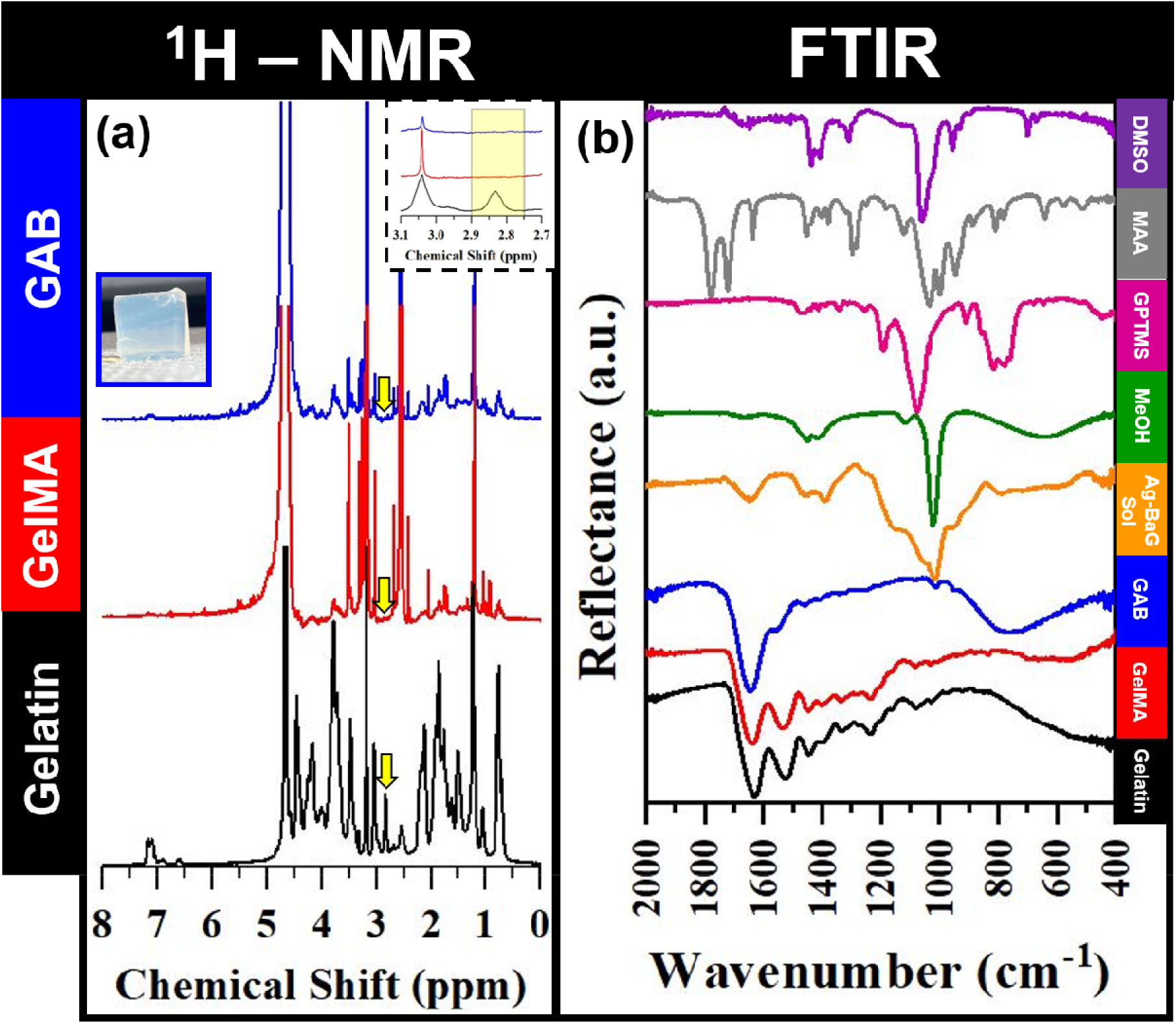
(a) ^1^H – NMR spectra of gelatin (as-received), GelMA, and GAB. Optical image inserts are additionally included to show the status of the hydrogels post-synthesis. The yellow arrows 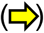 denote the lysine methylene signals used to determine the degree of substitution. A zoomed in spectra within this range is additionally shown as an insert for clarity. (b) FTIR spectra of the gelatin (as-received), GelMA, GAB, the Ag-BaG solution used during the GAB synthesis, methanol (MeOH), the coupling agent (3-Glycidyloxypropyl)trimethoxysilane (GPTMS), methacrylic anhydride (MAA) used for the methacrylation of the as-received gelatin, and dimethyl sulfoxide (DMSO) used as the solvent for the synthesis of GelMA and GAB.

Traditionally, GelMA has been synthesized in an aqueous environment such as PBS [63] and using methacrylic anhydride (MAA) as the methacrylation agent, leading to a maximum achievable DS of ~80 – 85% with large batch-to-batch variations. This was found to be a consequence of using an aqueous environment for the GelMA synthesis as the methacrylic anhydride preferentially hydrolyzes with its aqueous environment to form methacrylic acid, which is non-reactive with gelatin. To overcome this and achieve a DS of ~80 – 85% required using a 10 – 32-fold molar excess of MAA, compared to the lysine groups present in the gelatin [64]. Lengthy dialysis (> 7d) process was needed to remove the remaining cytotoxic reagents (i.e. MAA). Recently, it has been demonstrated that the use of a carbonate-bicarbonate system tuned to have a pH of 9.0 found success in increasing the DS up to 97% while dramatically lowering the MAA required to a 2.2 molar excess [64], however, a lengthy dialysis post-processing step was still required.

Here, we aimed to remove the adverse effects created by synthesizing GelMA in an aqueous environment through the use of an organic solvent, DMSO. It was hypothesized that the use of DMSO would prevent MAA hydrolysis and allow for ~100% DS to be achieved during synthesis. Furthermore, it was recently found that the dialysis step could be circumvented when using toluene as a precipitating agent [59]. The small dielectric constant for toluene (ε ~2.4), when exposed to the GelMA solution, increases the attractive forces between the oppositely charged portions of the GelMA allowing for agglomerations to grow resulting in the precipitation of the GelMA. An additional benefit to using the toluene precipitation method is that the DMSO and MAA are soluble in toluene allowing for their removal without the need for a dialysis process.

To verify the effectiveness of the toluene precipitation method, FTIR spectra (**Fig. 2b**) of gelatin as-received, GelMA, and GAB were collected in addition to the reagents used during the synthesis process of GelMA and GAB. As shown in **Figure 2b**, minimal spectroscopic changes were observed after the synthesis of GelMA; an indication that the precipitation method used was successful at extracting the GelMA from the solution. Additionally, the characteristic spectroscopic peaks for MAA and DMSO were not observed in the FTIR spectrum of GelMA (**Fig. 2b**) providing further evidence that the precipitation method was successful. For the GAB, the FTIR spectrum (**Fig. 2b**) presented the characteristic amide I, amide II, and amide III groups at ~1650 cm^−1^, ~1500 cm^−1^, and ~1450 cm^−1^ respectively [60] demonstrating the GelMA structure was preserved during the GAB synthesis, as expected. The broad Si-O bending peak ~750 – 800 cm^−1^ [60] is characteristic of the Ag-BaG indicative that there was successful incorporation and preservation of Ag-BaG during precipitation.

Given the successful synthesis of GelMA and GAB, when examining the morphology of both on the microscopic scale (**Fig. 3a-b, g-h**), clear morphological differences were observed. The GAB (**Fig. 3g-h**) showed thicker filaments of material roughly one micron in thickness with relative uniformity likely a result of the combination of GelMA and Ag-BaG. Interestingly, the combination of the Ag-BaG with GelMA led to a smoother surface morphology of the filaments compared to GelMA alone, where more node-like features were observed. The pore size of the GAB (**Fig. 3h**) was additionally present on the micron-scale, whereas the pores for GelMA (**Fig. 3b**) were evident on the sub-micron scale as a result of the sub-micron thick filaments.

**Figure 3.**
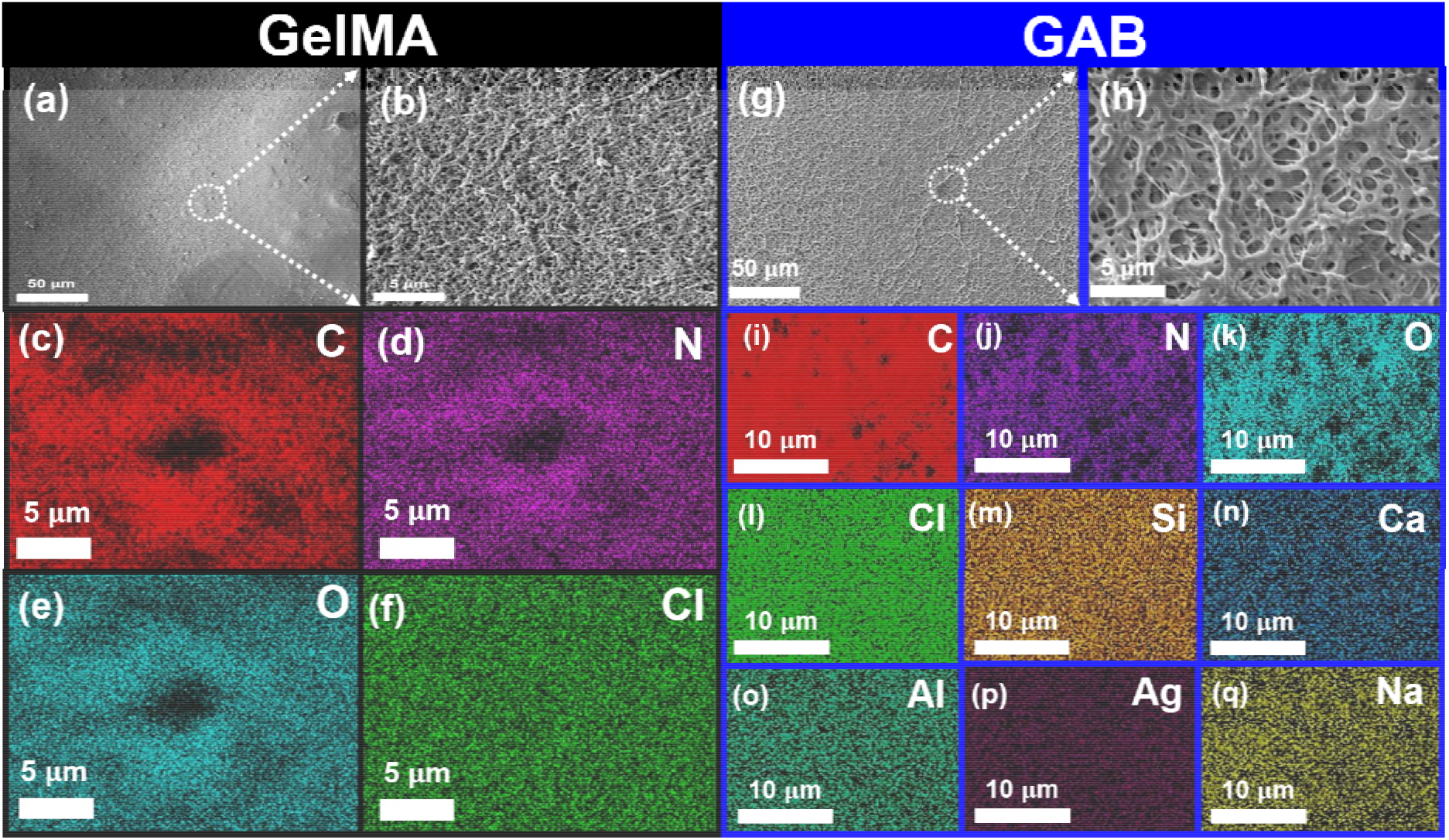
SEM images of (a, b) GelMA in addition to the respective EDS X-ray mapping, where (c) C, (d) N, (e) O, and (f) Cl were all found to be homogenously distributed down to the micron level. Additionally, SEM images of (g, h) GAB and corresponding EDS X-Ray mapping showing that (i) C, (j) N, (k) O, (l) Cl, (m) Si, (n) Ca, (o) Al, (p) Ag, and (q) Na are all homogeneously distributed down to the micron-level.

Regarding elemental homogeneity, both GelMA (**Fig. 3c-f**) and GAB (**Fig. 3i-q**) were found to have all elements homogenously distributed down to the micron level. This provides supporting evidence that the synthesis used to deliver GAB was additionally successful at achieving a high degree of homogeneity. This was critical as this provides supporting evidence that GAB should give a homogenous response when studied *in vivo*; allowing for the benefits of the GelMA and Ag-BaG to be exhibited simultaneously.

Regarding the performance aspects of GelMA and GAB in addition to the GelMA-Ag-BaG nanocomposite, the swelling ratio (**Fig. 4a**), pH evolution (**Fig. 4b**), and mass loss (**Fig. 4c**) were studied. It was found that the GAB underwent the least amount of swelling compared to the GelMA or the nanocomposite, although the difference in the swelling ratio between the nanocomposite and GAB was insignificant. The presence of the Ag-BaG in both the nanocomposite and GAB decreased the ability of the material to swell given the Ag-BaG does not experience swelling when exposed to aqueous environments. The swelling ratio (**Fig. 4a**) for GAB was likely decreased compared to the nanocomposite as a result of the covalent bonding that existed between the Ag-BaG and GelMA as opposed to the weaker Van der Waals interactions between Ag-BaG NPs and GelMA.

**Figure 4.**
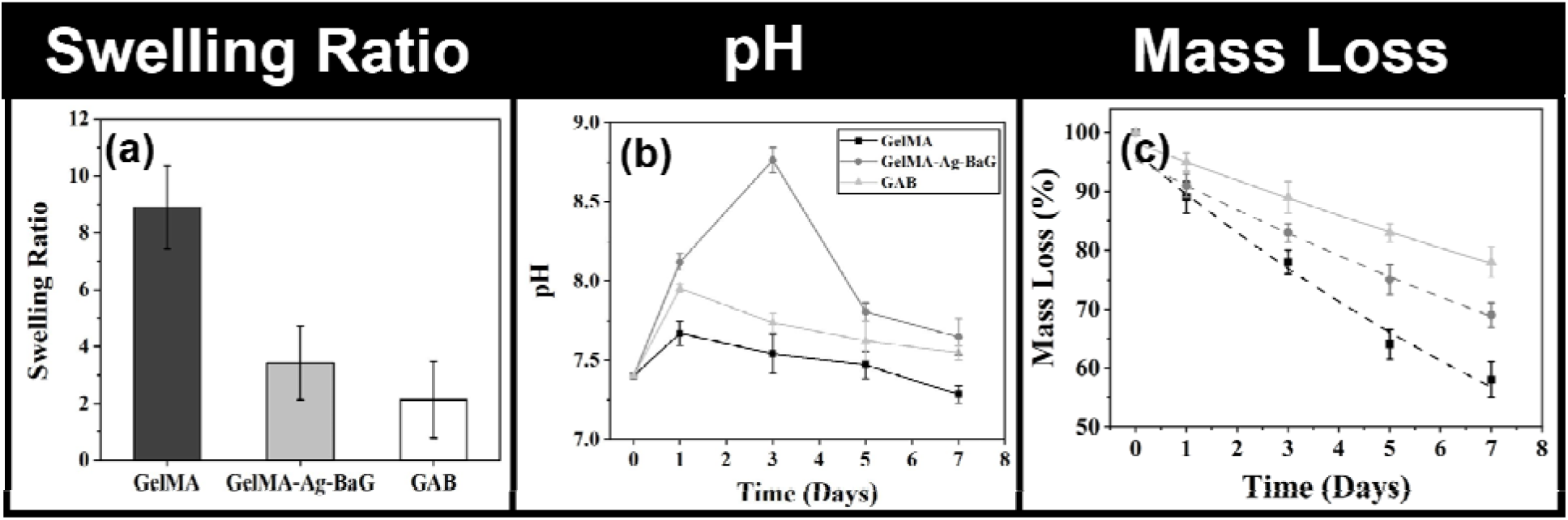
(a) The swelling ratio of GelMA, the GelMA-Ag-BaG nanocomposite, and GAB. (b) The pH evolution of GelMA, the GelMA-Ag-BaG nanocomposite, and GAB after 1, 3, 5, and 7d of immersion in PBS in addition to the corresponding (c) mass loss for each time point.

The pH evolution, studied over the course of seven days (**Fig. 4b**), revealed that GAB presented an evolution that modeled, well, the pH evolution of GelMA. This was likely due to the covalent bonding present in addition to the fine degree of homogeneity achieved during the synthesis process allowing the GelMA to mediate the ion release from the Ag-BaG compared to the nanocomposite. The GelMA-Ag-BaG nanocomposite displayed a large increase in pH that peaked ~8.8 before dropping below a pH of 8 after 5d of immersion. Sol-gel-derived bioactive glasses are known to have a burst release of ions [61] at early time points of immersion creating an alkaline environment, as evidenced in **Figure 4b**. This presents supporting evidence of the benefits that can be achieved when combining the GelMA with the Ag-BaG to create the hybrid material. Additionally, the GAB was found to be the most stable when the mass loss was studied up to seven days compared to the nanocomposite demonstrating further the advantages of covalently bonding the GelMA with the Ag-BaG.

To determine whether GAB was a viable material for 3D printing, its rheological performance was studied. **Table 1** shows a summary of the viscosity of GAB as a function of shear rate in addition to the storage and loss modulus and compared to values for GelMA obtained from literature along with the recommended values for each materials property for targeting 3D printing applications.

**Table 1.** Experimental and reported values for the rheological properties of GelMA-containing bioinks.

| Property | GAB Bioink<br>10% (w/v) GelMA |  |  | GelMA Bioink<br>10% (w/v)<br>Literature |  |  | Recommended<br>Values from<br>Literature |
| --- | --- | --- | --- | --- | --- | --- | --- |
| Viscosity (Pa·s) | 40,000 | ~1,000 | ~50 | ~300 | ~170 | ~40 | 0.03 – 60,000 |
| Shear Rate (s<sup>-1</sup>) | 0.1 | 1 | 10 | 0.1 | 1 | 10 |  |
| Storage<br>Modulus (Pa) | ~75,000 |  |  | ~800 – 1,000 |  |  | 100 – 1,000 |
| Loss Modulus<br>(Pa) | ~20,000 |  |  | ~5 – 40 |  |  |  |
| Shear Thinning | ✓✓✓ |  |  | ✓✓✓ |  |  | ✓✓✓ |

For viscoelastic materials such as GelMA and GAB, the storage modulus represents the ability of the material to absorb deformation energy elastically whereas the loss modulus represents the energy dissipated when a force is removed. For GAB, the storage modulus was found to be two orders of magnitude greater than either the GelMA reported in the literature or the recommended values (**Table 1**). The covalent coupling between GelMA and Ag-BaG allowed the material to absorb a much greater amount of deformation energy elastically by providing avenues where the strong ionic bonds from the Ag-BaG can contribute to the deformation energy storage. The loss modulus for the GAB was expectedly increased compared to GelMA given the presence of the Ag-BaG naturally increases the stiffness as a result of the orders of magnitude higher stiffness for bioactive glasses such as Ag-BaG compared to a softer material, such as GelMA.

The viscosity of the GAB (**Fig. 5b**) was found to decrease as a function of increasing shear rate, an indication of shear-thinning behavior. It is preferential for the GAB to exhibit shear-thinning behavior as shear thinning allows materials to exhibit self-healing abilities after removal of the shear forces allowing for higher quality constructs to be 3D printed. The linear region present between ~1 – 10 s^−1^ (**Fig. 5b**) was used to determine the flow consistency index (K) and flow behavior index (n) using a power fitting performed following equation 5.

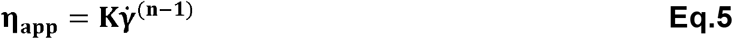

**Figure 5.**
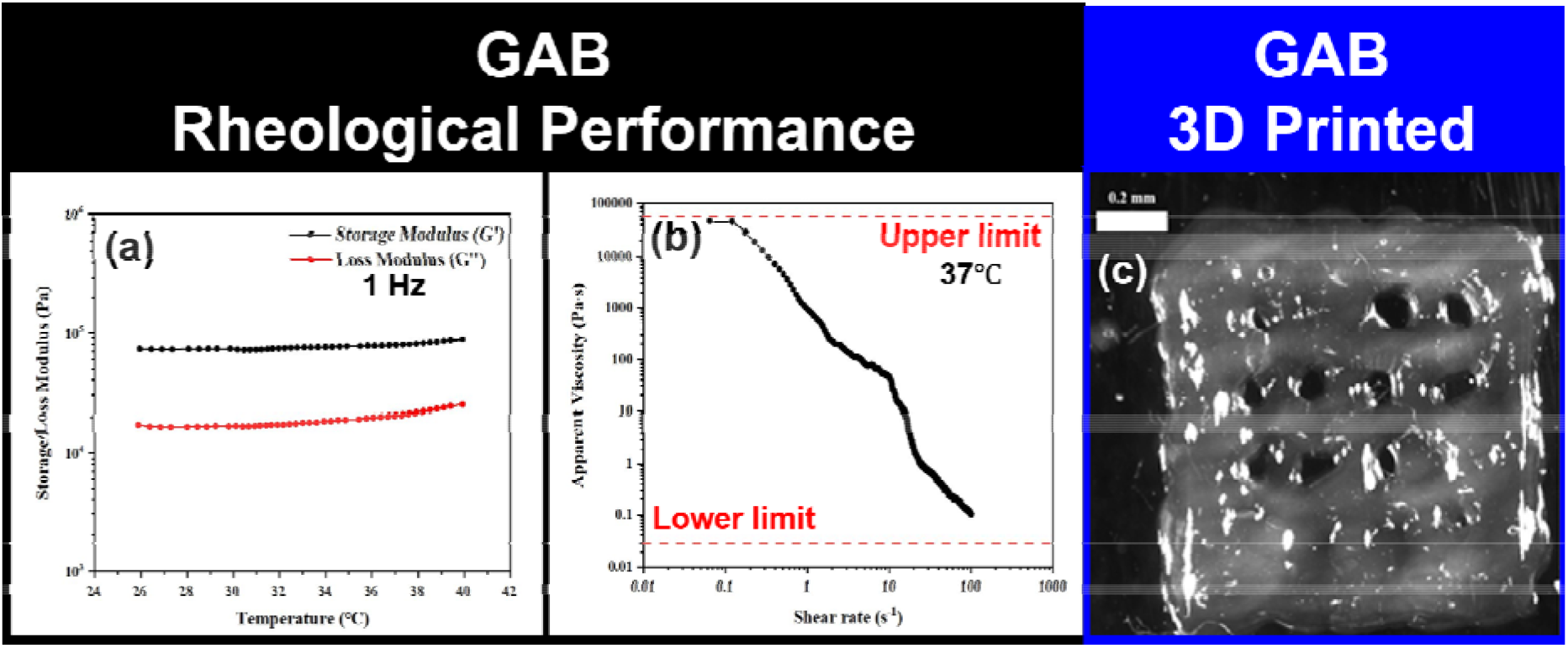
(a) The storage (G’) and loss (G”) modulus for GAB as a function of temperature ranging from 25°C to 40°C at 1 Hz of oscillation. (b) The apparent viscosity of the GAB as a function of the shear rate showing GAB exhibits shear thinning behavior. (c) A single mesh layer of GAB 3D printed.

From equation 5, GAB was found to have a flow consistency index (K) of 417±10 (Pa·s^n^) and a flow index behavior of 0.045±0.007 with a correlation coefficient (R^2^) of 0.98. Given the low value of n for GAB, this is further supporting evidence to suggest that GAB exhibits shear thinning behavior and furthermore behaves as a non-Newtonian fluid.

**Figure 5** showed a single-layer mesh 3D printed using GAB, where GAB was successfully printed using a pressure of 15 psi, an extruder temperature of 27°C, a 27 gauge metal tapered-tip, and a printing velocity of 4 mm s^−1^. The ability to 3D print GAB at 15 psi with an extruder temperature only 10°C cooler than physiological temperatures should minimize the shear forces and minimize the temperature differential cells would exhibit when incorporated into the material for 3D bioprinting. This presents supporting evidence of the printability of GAB and demonstrates its potential advantages for 3D bioprinting compared to other hydrogels such as GelMA. Furthermore, the ability to successfully incorporate Ag-BaG into GelMA without requiring the use of NPs is expected to improve cell viability during 3D printing given the introduction of additional shear forces due to the presence of the NPs.

## 3. CONCLUSIONS

Following our novel approach, chemically bonded GelMA-Ag-BaG (GAB) were successfully synthesized and cross-linkable enabling 3DP. The rheological evaluation found the GAB exhibited shear thinning behavior, which is a preferential characteristic for printability. The incorporation of the Ag-BaG was found to be homogenous at the molecular level that led the GAB to exhibit the least amount of swelling and the slowest degradation behavior compared to either the GelMA alone or the GelMA-Ag-BaG NP nanocomposites. GAB is expected to be suitable for extrusion-based 3DBP technologies and expected to improve cell viability of the 3D printed constructs as a result of the improvements in the characteristics of the material of the GAB bioink over a GelMA-Ag-BaG NP nanocomposite bioink.

